# Computational investigation of visually guided learning of spatially aligned auditory maps in the colliculus

**DOI:** 10.1101/2020.02.03.931642

**Authors:** Timo Oess, Marc O. Ernst, Heiko Neumann

**Affiliations:** Applied Cognitive Psychology, Ulm University, D-89081 Ulm, Germany; Institute of Neural Information Processing, Ulm University, D-89081 Ulm, Germany

## Abstract

The development of spatially registered auditory maps in the external nucleus of the inferior colliculus in young owls and their maintenance in adult animals is visually guided and evolves dynamically. To investigate the underlying neural mechanisms of this process, we developed a model of stabilized neoHebbian correlative learning which is augmented by an eligibility signal and a temporal trace of activations. This 3-component learning algorithm facilitates stable, yet flexible, formation of spatially registered auditory space maps composed of conductance-based topographically organized neural units. Spatially aligned maps are learned for visual and auditory input stimuli that arrive in temporal and spatial registration. The reliability of visual sensory inputs can be used to regulate the learning rate in the form of an eligibility trace. We show that by shifting visual sensory inputs at the onset of learning the topography of auditory space maps is shifted accordingly. Simulation results explain why a shift of auditory maps in mature animals is possible only if corrections are induced in small steps. We conclude that learning spatially aligned auditory maps is flexibly controlled by reliable visual sensory neurons and can be formalized by a biological plausible unsupervised learning mechanism.

## INTRODUCTION

Identifying a location of a visual or auditory event in our environment is advantageous for an organism to orient in space. Localizing a visual stimulus is a rather easy task, since the receptor array (sensorial neurons on a sensory organ) is topographically ordered. Hence, the location of a stimulus can be directly read out by its position on the retina. However, in the auditory domain localizing a stimulus is a complicated task that involves intensive computational steps to overcome two major obstacles. The challenge in sound source localization begins at the stage of the cochlea. The tonotopically organized receptor array represents neighboring frequencies but not adjacent spatial locations. Consequently, the location of a sound source cannot be directly inferred from its array position but needs to be computed using cues created by the head shadow, the distance between the ears or their shape. However, these cues lack associations to absolute locations in space. In order to establish such associations the brain utilizes vision as a guidance signal (Knudsen and Knudsen, 1989). Whenever a visual stimulus appears in temporal registration with an auditory stimulus, an association is established between the spatial location of the visual stimulus and computed auditory localization cues to form a topographically ordered representation of auditory sound source locations. Such associations can be found in the external subdivision of the inferior colliculus of barn owls in the form of a topographically aligned map of auditory space (Knudsen and Konishi, 1978).

Here, we present a neural model of auditory map formation in owls utilizing an unsupervised learning rule that evaluates correlations of sensory input streams. Map structures are defined as 1-dimensional arrays of topographically arranged single compartment neurons. Activations of model neurons are defined by rate-based changes of first-order conductance-based membrane dynamics and their transformation into firing rates. The model architecture is based on physiological findings in barn owls (Knudsen and Brainard, 1991; Hyde and Knudsen, 2000; Linkenhoker and Knudsen, 2002) and incorporates parts of external nucleus of the inferior colliculus (ICx) and the optic tectum (OT) to represent auditory and visual inputs, respectively. Unsupervised learning is defined by a neoHebbian 3-factor rule that incorporates an eligibility control signal, a co-activation plasticity mechanism, and a temporal trace of post synaptic activation (Gerstner *et al*., 2018). Together, these components enable stable map alignment of auditory space instructed by visual guidance signals.

Simulation results demonstrate the ability of the model to resemble behavioral and physiological findings in barn owls and explain the difference in remapping of auditory space in juvenile and adult owls when their visual field is shifted. Specifically, we show that the ability of remapping critically depends on the receptive field size of auditory neurons. This explains why a gradually induced prismatic shift enables remapping of auditory space in mature animals whereas a single large shift only works for juvenile owls.

## METHODS

The alignment of auditory space maps in barn owls occurs at the level of the midbrain between the central nucleus of the inferior colliculus (ICC) and ICx. This alignment is guided by retinotopic visual inputs from the OT. The ICC comprises tonotopically ordered neurons that are responsive to certain values of localization cues such as interaural time or level differences. By combining these cues over different frequency channels and associating them with visual information provided by the OT, the ICx forms a topographical map of auditory space.

Inputs to ICx model neurons at location *j* arise from ICC (audio) and OT (vision) and are denoted 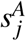 and 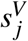, respectively. Each input is a one-dimensional vector of *N* = 40 entries that describes the input conductances to the ICx neuron population of size *N*. This number is chosen to achieve high input resolution while keeping the computational cost in a feasible range. Visual inputs from the OT are topographically structured whereas inputs from the ICC are tonotopical but show a topographical organization of localization cues (Feldman and Knudsen, 1997). Therefore, independent auditory 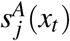 and visual 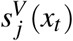 sensory input at location *j* can be modeled by

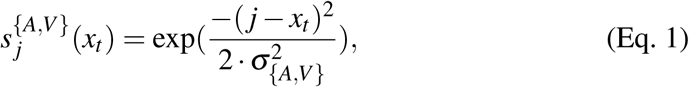

where *x*_*t*_ is the location at time *t* and *σ*_{*A,V*}_ the spatial extent of an auditory and visual stimulus on the receptor array, respectively. To ensure similar input strengths the input vectors are min-max normalized. An input location *x* is randomly chosen between 0 and *N* and for each such a location a Wiener process (initial setting: *σ*_*W P*_ = 1,*µ* = *x*) is applied that randomly selects five locations in the vicinity of *x*. Finally, these five locations are consecutively presented, each for 100 time steps (sufficient time to reach a steady state of the membrane potential), before choosing another initial location *x*. This process is repeated *N* times.

One model assumption is an increasing energy level in the visual map of space over time due to maturation of the visual system. This incremental increase of energy of visual inputs is modeled by filtering the visual input signal with a temporal high-pass filter according to 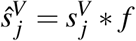 (*\**: convolution operator), where

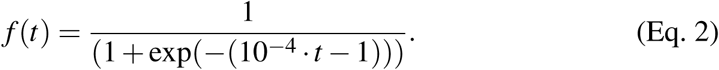

This leads to increased plasticity of the learning for more reliable visual input signals. The auditory input signal is fed to an ICx model neuron *r* _*j*_, whose membrane potential change is defined by:

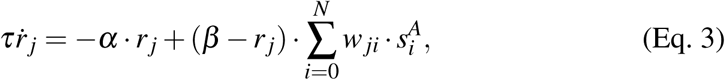

where 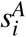 describes the auditory input, weighted by connection weight *w*_*ji*_ over all input locations. Parameter *τ* = 0.1 defines the membrane time constant, *α* = 1.0 is a passive membrane leakage rate and *β* = 1.0 describes a saturation level of excitatory inputs (standard neuron parameters). To create a firing rate, membrane potential *r* _*j*_ at time *t* is transformed by an output function *g*(*r*_*j*_(*t*)) = [*r* _*j*_(*t*)]_+_ = *max*(*r*_*j*_(*t*), 0). Note, that visual inputs do not drive the neuron auditory map neurons in accordance with neurophysiological findings (Knudsen and Knudsen, 1989).

The essential part of the model is a 3-component neoHebbian learning algorithm (Gerstner *et al*., 2018), that facilitates learning for spatially and temporally aligned inputs. Empirical exploration was used to choose best learning parameter values.

Weights are initialized with a large receptive field kernel, 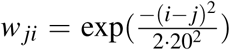 so that each receptive field ranges over the entire input space to replicate juvenile owls’ receptive fields. The weight adaptation Δ*w* for each learning step is governed by the 3-component structure

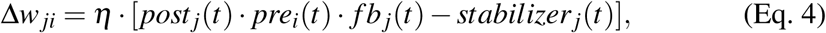

with *η* = 0.005 denoting the learning rate, 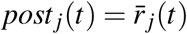 is the temporal trace value of the map activity 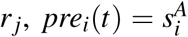 is the activity of the auditory input neuron and 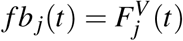 is an eligibility signal. This eligibility signal is driven by the activity of a spatially coincident visual neuron and the overall energy in the visual map, 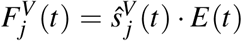 where is *E*(*t*) is the normalized total energy in the visual map and is calculated by 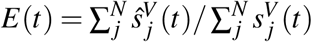. Here, 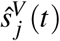 is the current activation of the visual input after filtering (see Eq. 2) and 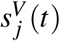 the maximal input strength before filtering. This mechanism guarantees that learning takes place only for reliable visual signals and for spatially aligned auditory and visual events (Brainard and Knudsen, 1993). The temporal trace value 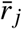 is calculated by 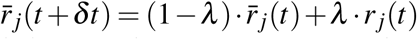 to achieve more robust map formation. *λ* = 0.5 denotes the trace parameter that defines the influence of previous values of *r* _*j*_ on its current state 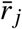.

The term *stabilizer*_*j*_(*t*) is added to counterbalance the correlative 3-component learning and is defined by 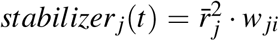 to pull weights towards constant energy by a rate proportional to the energy of the map neuron activation. By collecting all terms in the weight adaptation mechanism we arrive at a modified Oja learning rule (Oja, 1989) that incorporates an eligibility control signal to regulate the map learning process.

Connections from one neuron to another can degrade over time which is modeled by a decay function that is applied in each time step: *w*_*ji*_(*t* + *δt*) = (1 10*−*6) *w*_*ji*_(*t*). Here, *w*_*ji*_(*t*) can become very small but never 0. However, it is possible that connections eventually vanish completely. We model this process by *w*_*ji*_ = 0, for *w*_*ji*_ *<* 0.01. To compensate for this degeneration the model is endowed with a process that allows for reestablishment of already vanished weights. This is achieved by increasing weight values randomly in close vicinity to weights which values exceed a given threshold:

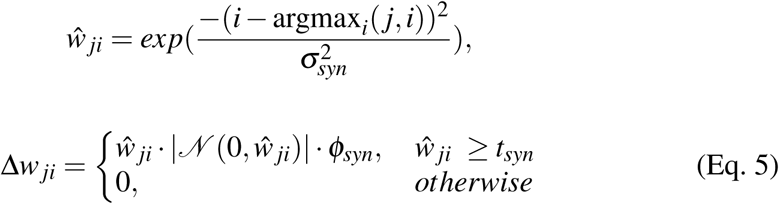

where *σ*_*syn*_ = 2.8 defines the range in which new weights can be created, *t*_*syn*_ = 0.5 is a threshold a weight has to exceed to initiate the process and *φ*_*syn*_ = 0.1 defines a scaling factor of the noise. This process is repeated every 1000th time step.

## RESULTS

In the following, we present simulation results that, first, show learning abilities of our model for spatially and temporally correctly aligned visual and auditory inputs. This learning of normal responses is used in a second experiment as a reference for comparison with learning for shifted visual inputs. In a third experiment, we show how the regained ability of shifting an auditory map for incrementally shifted visual inputs in adult owls depends on receptive field size of auditory neurons.

At the beginning of each simulation experiment the auditory system is in its juvenile state (reduced energy in visual map, broad receptive fields of auditory neurons, no auditory map alignment) and develops over the course of 50, 000 time steps to its mature state (maximal energy in visual map, narrow receptive fields of auditory neurons, and supposedly aligned auditory map). To demonstrate correct functionality of the model, we present a learned auditory map for temporally aligned, non-shifted visual and auditory inputs (Fig. 1 **A**). It represents correct map alignment in healthy owls. The abscissa indicates the spatial offset of the alignment from the predicted normal. The predicted normal describes the location offset between auditory and visual signals and is 0*°* for a perfectly aligned auditory map (spatial coincidence). Data presented here is collected by repeatedly presenting different sound source locations and measuring the response of auditory map neurons.

**Fig. 1:**
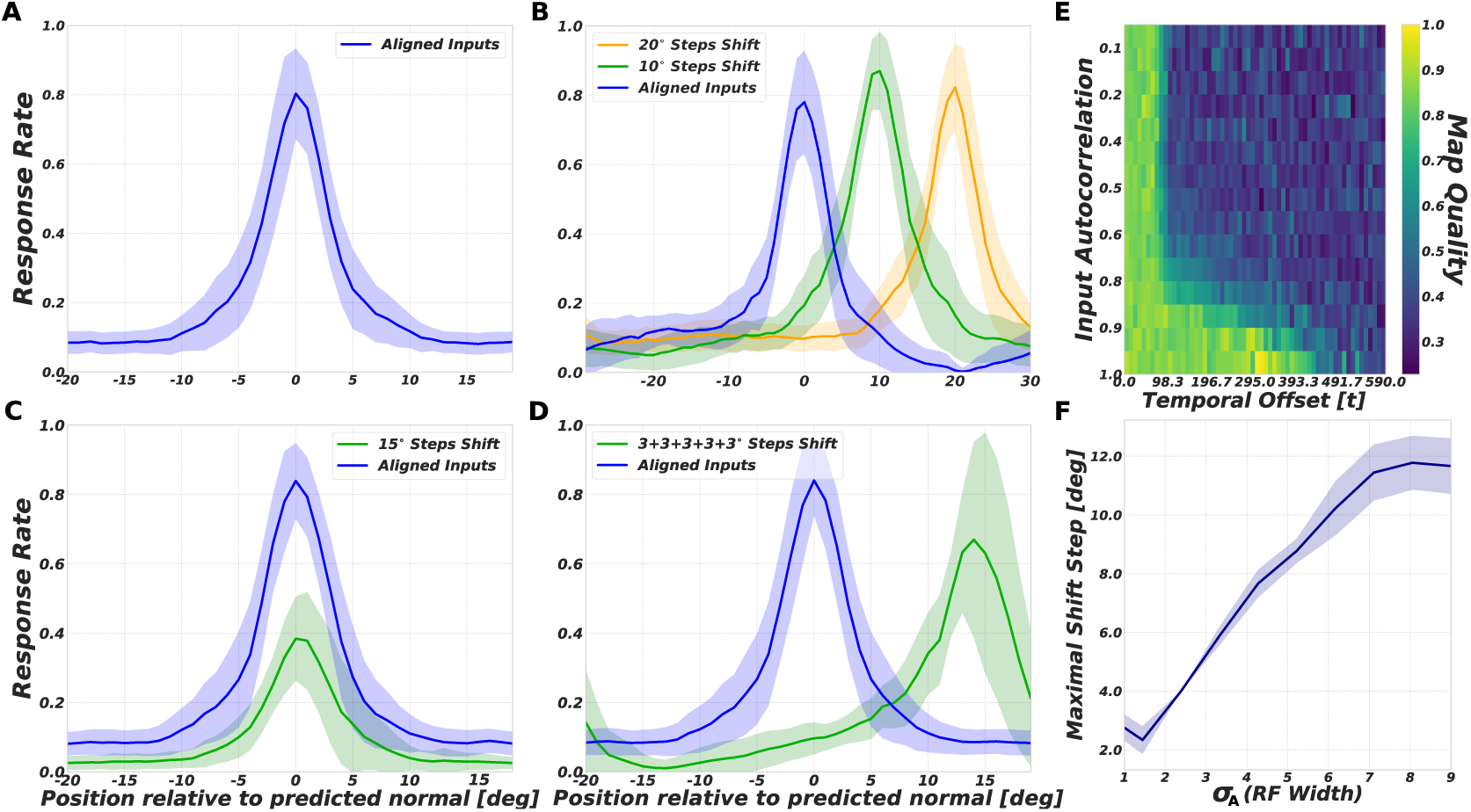
Left and middle column. Population tuning curves of ICx neurons to different auditory stimulus locations relative to each neuron’s predicted normal response are shown (as in Knudsen (1998)). Normal responses of a neuron are derived from the learned map for temporally aligned, non-shifted inputs. Lines depict mean of model neuron responses. Standard deviation in colored area. In all plots the blue line is the normal response plotted for easier comparison. **A** depicts model responses for temporally aligned, non-shifted inputs. **B** shows responses for various shifts of visual inputs. In **C** green line plots model responses for a shifted visual input of 15*°* in adult animals. **D** shows model responses for incremental shifted inputs in adult animals. Right column. **E** map alignment quality over autocorrelation value of inputs (ordinate) and their temporal offset (abscissa). Perfect map alignment is indicated in bright yellow, no alignment in dark blue. **F** depicts maximal shift over receptive field width when shift is introduced at a mature state.

Temporal coincidence between both inputs determines how well an auditory map can be aligned. For large temporal offsets the map alignment fails due to reduced activity of map neurons at the moment the visual signal would facilitate learning. Temporal coincidence is especially crucial if stimulus locations are randomly sampled from the environment (high *σ*_*W P*_ value of Wiener process). However, if consecutive auditory visual events are in spatial vicinity (they are spatially correlated, low *σ*_*W P*_ value) correct map alignment is still possible even for large offsets (Fig. 1 **E**).

Experiments with juvenile owls show that if a constant shift of visual inputs is introduced by prismatic goggles, the alignment of the auditory space map is shifted accordingly. This indicates a visually guided learning of auditory space (Brainard and Knudsen, 1993). We replicate this experiment by inducing a shift of the visual inputs by 10*°* or 20*°*, respectively, right at the beginning of the learning. Through its role as a guidance signal, the visually shifted input leads to a shifted alignment of the auditory space map (Fig. 1 **B**). However, this shift only occurs when introduced in juvenile owls, but remains ineffective when tested with adult owls (Linkenhoker and Knudsen, 2002). Our model exhibits the same behavior when presented with a non-shifted visual input during development (until time step 100, 000 to simulate sufficient adult experience) followed by a 15*°* shifted visual input. Map realignment fails since the shifted input is outside the receptive field range of auditory neurons (Fig. 1 **C**). Unlike the ineffective alignment modification in case of a single large shift, incremental small shifts in the visual input can lead to map realignment in adult owls. Simulations with multiple small incremental shifts of the visual signal demonstrate that our model is able to realign the auditory map of space in each consecutive step (Fig. 1 **D**).

Our results indicate that this phenomenon can be ascribed to the receptive field size of auditory map neurons. According to Hebbian learning, new connections and thereby realignment can only be learned if there is a temporally coincident pre- and post-synaptic activity of neurons (Hebb, 1949). If a large shift is introduced, the activity location of postsynaptic neurons in the auditory map and of visual input neurons do not coincide, since the stimulus location is outside of the spatially aligned auditory neuron’s receptive field. In contrast, if small shifts are introduced the stimulus location might still be in the receptive field range of the auditory neuron which leads to an activation and therefore a relearning of connections. We tested this hypothesis by measuring the maximal shift step size depending on the receptive field size of the auditory neurons. For each receptive field size (range [0.5 *· σ*_*A*_, 7 *· σ*_*A*_] *≈* [4*px*, 40*px*]) different shift steps (range [5*°*, 25*°*] = [5*px*, 25*px*]) are tested and the maximal possible shift is measured (a shift is successful if the activity location of the map neurons corresponds to the induced shift). In total, we ran 8 simulations and calculated the mean maximal shift for each receptive field size. For increasing receptive field size the maximal shift step becomes proportionally larger (Fig. 1 **F**).

## DISCUSSION

We presented a neuron model that learns aligned auditory maps of space by applying a 3-component neoHebbian correlative learning rule to its visual and auditory inputs. Previous investigations of auditory map alignment demonstrated that alignment is possible with a *map adaptation cue* in a spiking neuron model (Huo and Murray, 2009) or with a simple, unconstrained Hebbian learning rule for visual and auditory inputs (Witten *et al*., 2008). Our model differs in that it directly uses visual inputs as a guidance signal for map alignment and is able to explain the unaffected map alignment in adult owls when a large prismatic shift is induced versus the ability to realign for incremental shift step size.

Simulation results demonstrated the model’s ability to successfully learn aligned auditory maps of space for temporally and spatially aligned visual and auditory sensory inputs. Due to the introduced eligibility signal, visual inputs do not drive responses of auditory map neurons but merely elicit learning for coincident stimuli, which is important for close replication of biological findings.

The incorporated trace rule can compensate for small temporal offsets between the visual and auditory inputs. However, for large temporal offsets, results indicate that the success of learning strongly depends on the spatial autocorrelation of the input locations. That is, if auditory and visual inputs are not randomly sampled in space but are chosen according to a Wiener process with small *σ*_*W P*_, correct map alignment is still possible. This implies that learning is enhanced for stimulus locations that show strong autocorrelation. When transferring our results to real world scenarios, this enhancement is of special interest since audio-visual events rarely happen to occur at just a single location, followed by other random locations but are likely to happen in spatial vicinity. Therefore, such an enhanced learning capability for strongly autocorrelated input locations seems to facilitate learning of real world stimuli.

It has been shown experimentally that the ability to shift the auditory map alignment is reduced for adult animals (Linkenhoker and Knudsen, 2002). Our model results predict that the receptive field size of the auditory neurons is responsible for this reduction and it can serve as an index of maximal shift step size for visual inputs that can still induce map realignment. This prediction could be tested by varying the prismatic shift step size and determining the receptive field size for adolescents of different ages. If our prediction is correct the two values should correlate.

Despite the presented results, at the moment our model is incapable to maintain already established maps and to quickly readjust to normal vision as it has been demonstrated for owls in (Knudsen, 1998). However, we argue that this could be achieved by adapting the learning algorithm and extend the input by elevation cues. Together with a N-methyl-D-aspartate (NMDA) receptor signal which in addition to the eligibility signal could control the learning, we expect the model to further extend its capability to resemble neurophysiological and behavioral studies.

## Notes

### Competing Interest Statement

The authors have declared no competing interest.

### Summary of Updates

Minor changes for clarification

